# Beyond linear regression: mapping models in cognitive neuroscience should align with research goals

**DOI:** 10.1101/2021.04.02.438248

**Authors:** Anna A. Ivanova, Martin Schrimpf, Stefano Anzellotti, Noga Zaslavsky, Evelina Fedorenko, Leyla Isik

## Abstract

Many cognitive neuroscience studies use large feature sets to predict and interpret brain activity patterns. Feature sets take many forms, from human stimulus annotations to representations in deep neural networks. Of crucial importance in all these studies is the mapping model, which defines the space of possible relationships between features and neural data. Until recently, most encoding and decoding studies have used linear mapping models. Increasing availability of large datasets and computing resources has recently allowed some researchers to employ more flexible nonlinear mapping models instead; however, the question of whether nonlinear mapping models can yield meaningful scientific insights remains debated. Here, we discuss the choice of a mapping model in the context of three overarching desiderata: predictive accuracy, interpretability, and biological plausibility. We show that, contrary to popular intuition, these desiderata do not map cleanly onto the linear/nonlinear divide; instead, each desideratum can refer to multiple research goals, each of which imposes its own constraints on the mapping model. Moreover, we argue that, instead of categorically treating the mapping models as linear or nonlinear, we should instead aim to estimate the complexity of these models. We show that, in many cases, complexity provides a more accurate reflection of restrictions imposed by various research goals. Finally, we outline several complexity metrics that can be used to effectively evaluate mapping models.

## 1 INTRODUCTION

In recent decades, neuroscientists have witnessed a massive increase in the amount of available data, as well as in the computational power of the tools we can apply to the data. As a result, we can now leverage huge datasets to build powerful models of brain activity. In this era of new opportunities, it is important to be mindful of conceptual choices we make before modeling our data. This paper discusses one such choice: the choice of a mapping model that relates features of interest to brain responses.

When studying a brain circuit, area, or network, it is often useful to formulate and test hypotheses about features that elicit a response in the relevant neural units^1^ (a single cell, a population of neurons, a brain area, etc.). The features can be stimulus-based (**Figure 1A**), behavior-based (**Figure 1B**), or based on responses in other neural units within the same brain or in a brain of another individual (**Figure 1C**). The ways in which the features are derived vary widely: common sources include human annotations (e.g., “faces” and “scenes”), empirical measurements (e.g., behavioral or neural responses), and outputs of a computational model (e.g., a vector of responses to each image in a layer of a deep neural network (DNN); **Figure 1D**).

**FIGURE 1.**
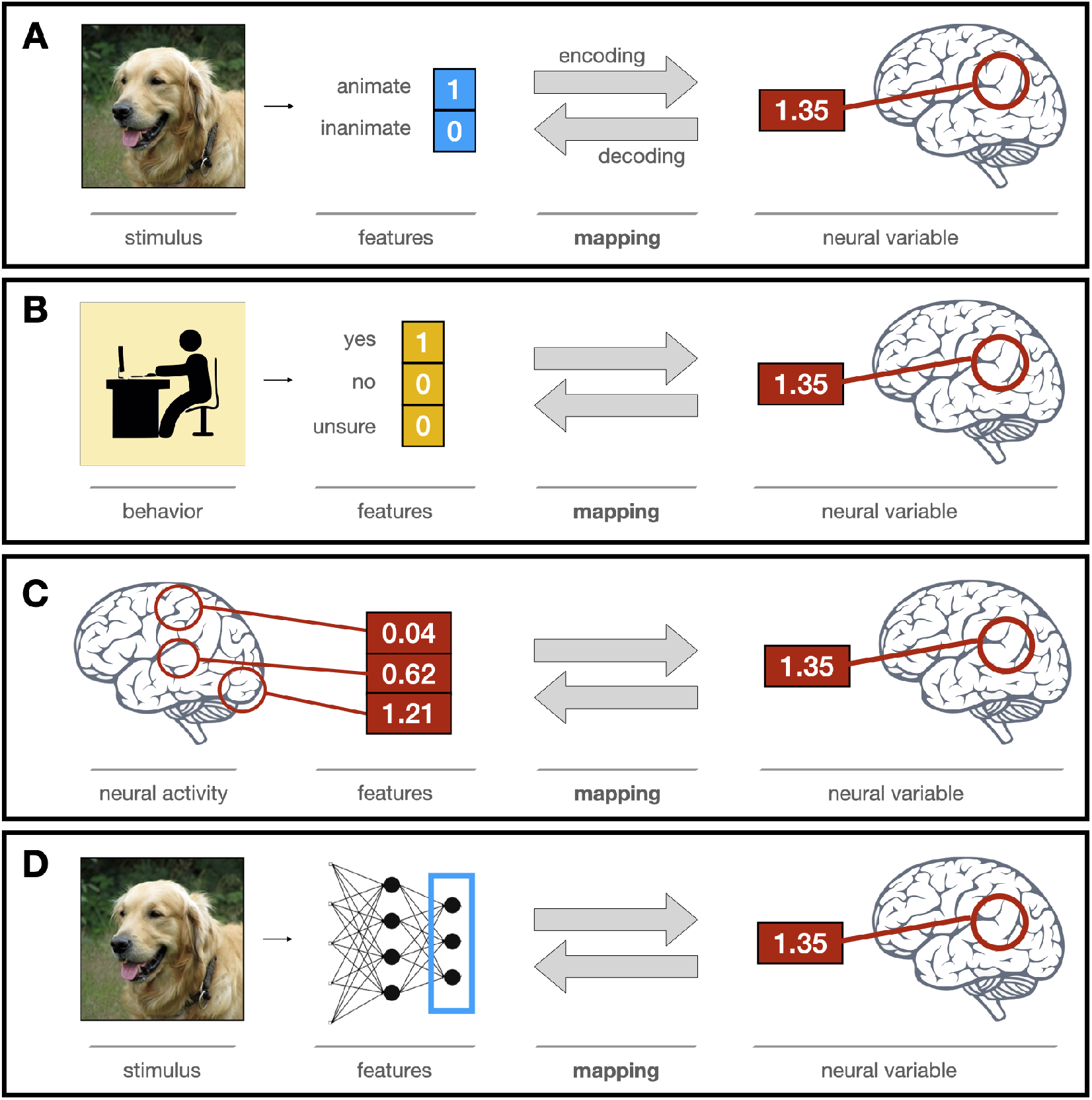
The encoding/decoding modeling framework in cognitive neuroscience. **(A)** Studies investigating the effect of external stimuli on brain activity start with the stimulus, extract its features of interest, and use a mapping model to establish the mapping between these features and a neural variable extracted from the data recorded during/after stimulus presentation. **(B)** In other studies, researchers extract features associated with participants’ behavior and map those onto the neural variable recorded before/during this behavior. **(C)** Another class of studies describes the mapping between activity in different brain regions where neural variables serve as the features. **(D)** In recent years, more and more studies replace hand-crafted features, like those shown in (A), with high-dimensional feature vectors derived from models of brain function, such as neural networks.

To relate a set of features to brain activity, we need to establish a mapping between them. The exact form of the mapping is typically learned from the data, although the space of possible mappings is defined in advance (e.g., linear functions).

Why is a mapping necessary? In principle, we could limit ourselves to mapping-free models, which use features of interest and a set of fixed parameters to predict neural activity directly. For instance, given some information about the stimulus, a mapping-free model would predict the exact firing rate of a particular neuron or change in BOLD activity in a given voxel. However, mapping-free models in cognitive neuroscience today are almost always infeasible. One limitation is a mismatch between the granularity of our theoretic predictions and the measurements to be modeled. For instance, we might want to test a model of brain function that predicts an increased firing rate in a neuron in response to a face but does not specify the exact amount of that increase. The second limitation is the lack of a priori knowledge about functional differences between neural units that anatomy fails to explain (e.g., which exact neurons will respond to faces in a given individual). Thus, we might want to test our predictions against the neural data without deciding a priori which neural units (or combinations whereof) might encode the features of interest. The third limitation is the data that we use, which typically provide a noisy and/or indirect measure of neural activity. The presence of noise and/or an underdetermined linking function between our predictions and our measurements means that we often want to incorporate some free parameters to infer this information from the data. All in all, when modeling neural data, some level of fitting is almost always required.

A mapping model is a model that relates features of interest and neural data^2^. Its main distinguishing feature is the presence of free parameters whose values are determined in the process of training the model on neural data. This makes mapping models meaningfully different from models of brain function, which aim to mimic neural computations but are not trained on neural data (**Figure 2)**. In principle, a model of brain function can also have its parameters trained or fine-tuned using neural data [5]; in this case, the distinction between the two becomes blurred. In practice, however, the majority of studies today separate these two steps: a model of brain function can be used to derive features of interest, and a mapping model is then fitted to link these features and neural data.

**FIGURE 2.**
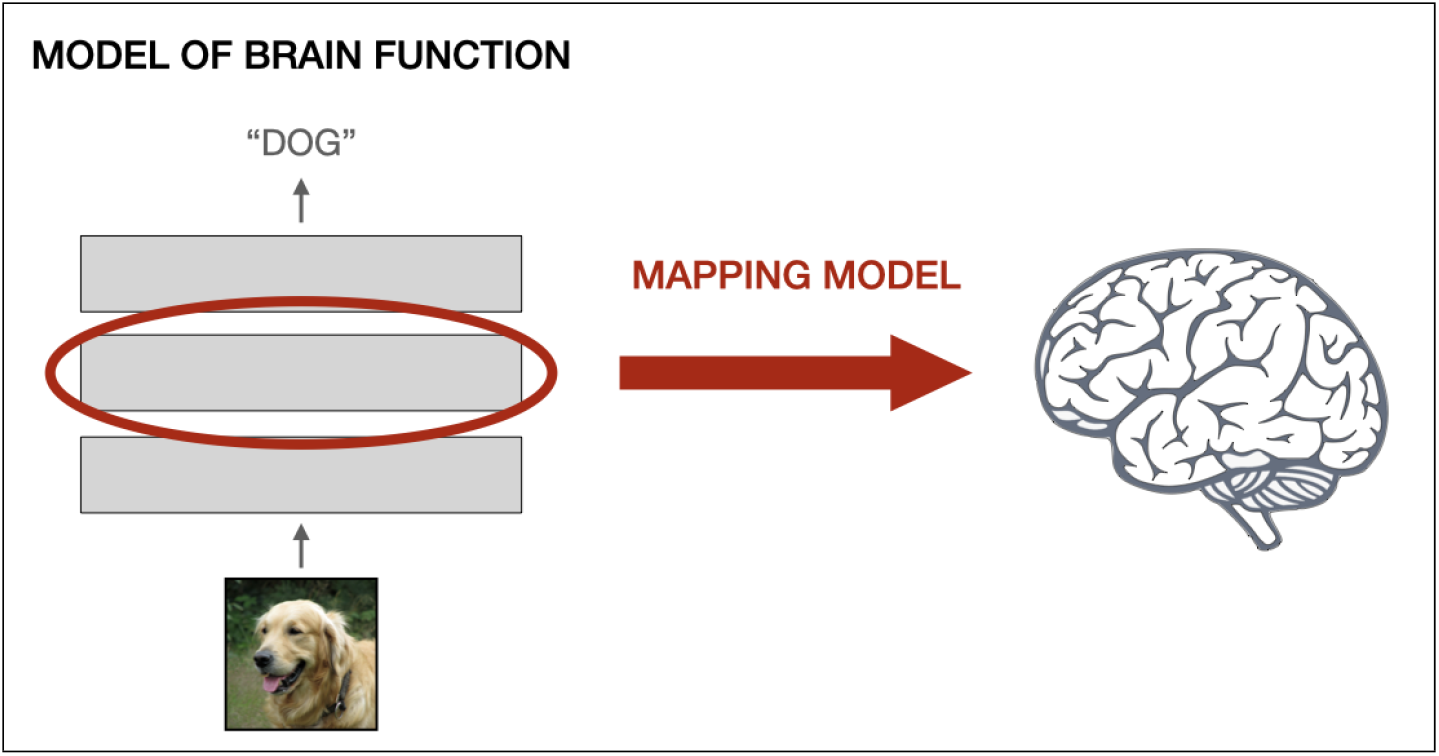
The distinction between a model of brain function and a mapping model. A model of brain function aims to mimic the brain, but does not directly map onto neural data. Our focus is on mapping models, which link a feature set to a neural variable. The mapping model depicted here uses features derived from a model of brain function (like in **Figure 1D**).

Mapping models have many properties that researchers need to take into account, but the most common distinction is drawn between (a) a linear mapping model (such as linear regression) and (b) a nonlinear mapping model (such as a neural network).

## 2 THE CONTROVERSY

Today, the vast majority of cognitive neuroscience studies use linear mapping models (such as linear regression). Linear mapping models are ubiquitous across domains (including vision, audition, and language) and recording techniques (from single unit recordings to non-invasive measures such as M/EEG and fMRI). Linear mapping models are often used in conjunction with nonlinear models of brain function, a common approach often referred to as a linearized model. In linearized models, a predefined nonlinearity is applied to a stimulus and/or a neural variable to derive a set of features, which are then linked to neural data using a linear mapping model. Linearized models were first established in sensory neuroscience [6] and have become the predominant approach, particularly in encoding models [4, 7, 8].

One of the major reasons that linear mapping models have become so common is a practical one: they can be fitted using standard statistical techniques that require little data and modest amounts of computation. However, recent advances in machine learning (ML), coupled with an increased availability of computing resources, have enabled neuroscientists to explore a much broader range of modeling techniques [9]. The concurrent increase in the size of available datasets [10, 11, 12] has enabled researchers to train large-scale mapping models without overfitting them. As a result, a number of applied neuroscience studies have leveraged the power of ML-based methods to build flexible nonlinear mapping models and use them to identify neural correlates of brain disorders [13, 14, 15, 16] and of behavioral traits [17, 18, 19].

Although these computational advances have substantially diminished the practical constraints on the mapping models (cf. **Section 4)**, many researchers still prefer to use linear mappings, arguing that such mappings are preferable not only for practical, but also for theoretical reasons. Some of the common theoretical arguments in favor of linear mapping models are the following:

1. Linear mapping models facilitate a comparison of predictive accuracy across feature sets [20, 21, 22, 23].
2. Linear mapping models estimate weights for individual features, making the mapping more interpretable [24, 25, 4, 26, 27, 3].
3. Linear mapping models are more biologically plausible: they approximate readout by a downstream area and can therefore indicate what information is available to the rest of the brain [28, 29].

However, none of the three general desiderata that are relevant to mapping model selection — predictive accuracy, interpretability, and biological plausibility — align straightforwardly with the linear/nonlinear mapping model choice. In the following section, we critically review these desiderata and show that each of them, in fact, can refer to several distinct research goals, each of which places its own demands on the space of mappings to be considered, meaning that the arguments above only apply in a subset of research scenarios and cannot serve as general guidelines.

## 3 HOW DO NEUROSCIENTISTS USE MAPPING MODELS?

To choose the best mapping model, we first need to specify the goal that we are trying to achieve [30, 31, 29]. The goal, of course, can be defined at various levels of granularity. Here, we focus on specific goals a researcher might want to accomplish by training a mapping model on their data (as opposed to high-level research goals, such as curing Alzheimer’s or building artificial general intelligence).

We categorize research goals under three general mapping model desiderata^3^ — predictive accuracy, interpretability, and biological plausibility. We show that each desideratum, in fact, corresponds to several distinct goals with their own mapping model requirements, and that the linearity requirement only holds for some of the goals.

### 3.1 Predictive accuracy

In neuroscience, as in other fields, scientific progress is driven by the generation of new hypotheses, followed by the testing of the hypotheses’ predictions against experimental data, and then by the selection of the best (most accurate) hypothesis (or generation of new hypotheses if none of the current hypotheses are good enough). In the encoding/decoding framework, a hypothesis can be operationalized as a set of features hand-crafted by the researchers, (e.g., [32]), derived from a model of brain function (e.g., [23]) or obtained from behavioral ratings (e.g., [24]). A common way to measure the predictive accuracy of a set of features is to use a mapping model that will estimate the best link between the features and the neural data. The mapping — fit on the training set — can then be used to predict responses in a held-out test set, after which we can evaluate those predictions by, e.g., correlating them with the test data. These correlations are often normalized by an estimate of the reliability of the data (a “ceiling”) to yield an estimate of explained variance [33, 34, 35, 36, 37, 38, 39, 40, 41].

This prediction-oriented framework can be used to achieve multiple research goals, only some of which impose specific constraints on the mapping model.

#### 3.1.1 Test feature relevance

One research question a neuroscientist might ask is “do neural data *Y* contain information about features *X*?” In this scenario, the goal is to find a mapping model that allows us to achieve significant (above-chance) predictive accuracy. Above-chance model performance can have both theoretical relevance (for instance, an encoding model showing that features *X* drive a certain brain region *Y*) and clinical relevance (for instance, a decoding model that predicts disease severity *X* based on neuroimaging data *Y*). Formulated this way, the above-chance-accuracy approach does not have to commit to the exact format of feature-brain mappings — all that matters is the mapping model’s performance on held-out data. For instance, a study that finds information about an imagined visual scene (*X*) in primary visual cortex [42] could in principle provide a valuable contribution to the field even if it used highly unconstrained nonlinear mappings. Similarly, if researchers aim to predict whether a given case of mild cognitive impairment will develop into Alzheimer’s within the next year, they do not have to pre-specify (or limit) the exact nature of the mappings they will consider. All in all, for studies in this category, the main objective is for the mapping model to achieve significant predictive accuracy (or, for applied research, to reach a certain accuracy threshold), and the space of possible mappings should be large enough to allow it.

#### 3.1.2 Compare feature sets

Predictive accuracy can be used to compare competing feature sets (often extracted from different models of brain function) with the goal of selecting the one that best fits neural data [43]. This goal is achieved by fitting encoding mapping models from several different feature sets to a pre-selected set of neural responses and comparing the variance explained by each mapping. A question that these studies tend to ask is: “Which feature set provides the most faithful reflection of the neural representational space?” When performing a systematic comparison of feature sets, researchers often choose to restrict themselves to linear mappings because a powerful non-linear mapping model could inadvertently incorporate transformations that reduce or erase the differences across feature sets. For example, if the goal is to determine whether activity in inferior temporal cortex is better predicted by an early or a late layer of a convolutional neural network, we should use a mapping model with a limited expressive power; otherwise, the mapping model will be able to transform features from an early layer into features from a late layer, eliminating meaningful differences between them. Thus, feature comparison studies often benefit from linear mapping models.

#### 3.1.3 Build maximally accurate models of brain data

Finally, some researchers are trying to build accurate encoding models that, in essence, enable simulations of neuroscience experiments [44, 45]. This type of modeling is especially important in cases when experimental data are expensive or hard to obtain: with a high-accuracy model of brain responses, a researcher can run thousands of experiments *in silico*, refine their hypothesis, and then test the critical predictions *in vivo*. Furthermore, these models may become an important component of testing experimental replicability: if the same phenomenon is shown both *in vivo* and *in silico*, it is less likely to be a false positive. In contrast to mapping models used for feature comparison (**Section 3.1.2)**, models used for *in silico* experiments are only useful if they clear a very high accuracy bar (ideally close to the noise ceiling of neural data); otherwise, experiment simulations they produce would not be trustworthy. The nature of the features used for these maximally accurate models is also irrelevant: rather than testing the performance of specific, pre-selected feature sets, the researcher can select any set that achieves the desired results, even if these features end up being raw picture pixels. The best way to build these *in silico* brains might be to train large powerful mapping models on large amounts of neural data. In this scenario, there is no theoretical justification for a linear mapping constraint because, as in **Section 3.1.1**, the primary goal is maximizing predictive accuracy on held-out data.

### 3.2 Interpretability

Once we find a mappingthat achieves sufficiently high predictiveaccuracy, we often want to interpret it. Which features contribute the most to neural activity? Do neurons/electrodes/voxels respond to single features or exhibit mixed selectivity? How does the mapping relate to other models or theories of brain function?

The traditional view is that linear mappings are easier to interpret than non-linear mappings [4]. However, the goal of building interpretable models is ultimately complicated by the fact that a clear-cut definition for interpretability is lacking. Below, we discuss three definitions of interpretability, ranging from strictest to loosest, and show that they provide different constraints on the mapping models. Importantly, in each of these cases, interpretability places restrictions not only on the mapping model, but also on the features that can be used to yield meaningful interpretations.

#### 3.2.1 Examine individual weights in the mapping model

Traditionally, many cognitive neuroscientists have aimed to interpret a neural signal by identifying a set of words to describe its function (as in [46, 32]). In this scenario, a useful model of brain activity has features that can be described using one or a few words (“faces”, “vertical lines”, etc.) — a property that is often referred to as nameability — and a linear mapping between these features and neural data. The dimensions of the neural data are also nameable (e.g., a brain voxel with certain coordinates or an electrode placed in a specific part of the brain), although in practice this criterion can be more loose. Under this setting, the weights of a linear mapping model can be interpreted as a relative measure of contribution of each input feature to each output feature (this can be done regardless of mapping direction).. We consider this to be the strictest definition of interpretability because it places the strongest constraints on both the features (which have to be nameable) and the space of possible mappings (which have to be linear).

Although this definition of interpretability is perhaps the most intuitive, it suffers from two shortcomings. First, weight interpretation is difficult in cases where the features (for an encoder) or the brain recordings (for a decoder) suffer from multicollinearity and/or when regularization is used to impose a prior on model weights [27, 3]. These issues are especially pervasive in the case of fMRI decoding models, where the brain measurements from nearby voxels are almost always correlated and where researchers often use regression with L2 regularization (also called ridge regression) to deal with the large number of voxels and noise in individual voxels. Second, feature nameability may be an overly restrictive metric, as it limits our understanding to a vocabulary that is heavily biased by *a priori* hypotheses and may not include words for the concepts we actually need [47]. For instance, recent work has shown that (a) neurons typically described as “face-responsive” respond more strongly to artificial images produced by DNNs than to natural images described by the word “face” [48]. and (b) a neural-network-based linearized model of activity in the fusiform face area predicts responses to faces better than label-based models [45], suggesting that simple verbal features cannot provide a full account of neural activity. To overcome the limitation of using individual nameable features, many researchers have instead started to use high-dimensional feature sets.

#### 3.2.2 Test correspondences between representational spaces

A looser notion of interpretability, which has become popular in the last decade, relies on the use of high-dimensional feature vectors that are linearly mapped, usually via encoding, to a neural variable [49, 23]. As mentioned in **Section 2**, this setup is commonly referred to as “linearized” models of brain function. When using large-scale feature sets, we cannot always interpret the weights of a linear mapping model in the same way as we did with nameable features. If individual features within a set cannot be labeled and/or are derived via a sequence of nonlinear operations (e.g., in the case of DNN layer activations), examining individual features has a limited potential to inform our intuition [30]. However, we can examine the feature set as a whole, to ask: do features *X*, generated by a known process, accurately describe the space of neural responses *Y*? Thus, the entire feature set becomes a new unit of interpretation, and the linearity restriction is placed primarily to limit the space of possible feature space transformations. For instance, the finding that convolutional neural networks and the ventral visual stream produce representational spaces that are similar up to a linear transformation (e.g., [23, 20, 50, 22]) allows us to infer that both processes are subject to similar optimization constraints [51]. That said, mapping models that relate two representational spaces do not have to be linear, as long as they correspond to a well-specified hypothesis about the relationship between them; for instance, we might want to relate the intrinsic dimensionality of the spaces being compared, an approach that is inherently nonlinear ([52, 53]; for discussion, see [54]).

#### 3.2.3 Describe the feature set as a whole

The loosest definition of interpretability is the ability to name and/or describe the set of features that was used to train the mapping model (e.g., “phonological features”). In this scenario, we make no assumptions about a particular representational geometry of these features (such as linear separability). The lack of specific assumptions about the form of the feature-to-brain mapping means that constraints on the mapping model are not strictly necessary — all we need is an epistemologically satisfying description of the features that would apply regardless of which mapping is applied to these features. If a mapping model that uses these features achieves good predictivity, we can say that a given set of features is reflected in the neural signal. Under this definition, any mapping model (encoding or decoding) is interpretable as long as we can describe the set of features that it uses.

### 3.3 Biological plausibility

In addition to predictive accuracy and interpretability-related considerations, biological plausibility can also be a factor in deciding on the space of acceptable feature-to-brain and brain-to-feature mappings. Given that the mapping model is commonly used to determine which features are reflected in neural data, it is important to select the space of mappings in a way that can lead to the selection of biologically plausible feature sets.

#### 3.3.1 Simulate linear readout

One of the main arguments in favor of linear mappings is the claim that they approximate the linear readout performed by a putative downstream brain area [28, 29]. Under this view, the mapping model approximates the transmission of the features to a hypothetical information consumer. The linear readout requirement often serves as a proxy for feature usability: if the features can be extracted with a linear mapping model, it means that they require few additional computations in order to be used downstream. Note that this argument only applies to decoding mapping models, which match the direction of the readout (from the brain area to a hypothetical consumer).

The ability to use features of interest in downstream computations is indeed an important consideration. However, there are reasons to be cautious about the linear readout requirement. First, some models operate on neural data that are collected from multiple recording sites rather than a single neural population/region, which makes subsequent linear readout biologically implausible. For instance, decoding models that use whole-brain data, such as M/EEG, have no downstream region that could ‘read out’ information from all over the brain — the only entity performing readout is the observer. Second, linear readout might not be an accurate characterization of the decoding mechanisms used by downstream areas to extract information from the brain region of interest. In fact, unlike linear models that pool across all measured neurons or voxels in the region of interest, readout in biological neural systems is likely to be both sparse (e.g., [55, 56, 57, 58]) and nonlinear [59, 60, 61, 62, 63]. Third, linear regression is a fairly arbitrary threshold to draw for mechanistic plausibility. A linear mapping model can extract many features from the data, some of which do not faithfully reflect the underlying neural computations and could not possibly be read out by a downstream neuron. For instance, fMRI signals from V1 contain voxel-level biases that allow orientation decoding (such as radial biases in the retinotopic map) that are distinct from orientation-related neural computations (such as activity in orientation-specific cortical columns), which results in a mismatch between information used by the mapping model and information used by actual neurons [64]. In sum, unconstrained linear mapping models (or linear mapping models constrained by weight distribution among many features, like ridge regression) may be both overly limiting because they do not account for possible nonlinear computations and overly greedy because they might leverage information in a way that real neurons do not.

Is there a better mapping model that accounts for possible nonlinear computations during readout without being overly broad? One approach is to introduce parsimony constraints on the feature spaces [65]. For instance, introducing a sparsity constraint (i.e., allowing the mapping model to access only a limited number of neurons) could increase the biological plausibility of putative readout [66]. However, in the context of measurements that collapse across large numbers of neurons (i.e., most measurements in cognitive neuroscience), the sparsity constraint might be impossible to enforce, as a single voxel or electrode already combines signal from a large number of neurons. More broadly, evaluating the biological plausibility of decoding is difficult as readout might differ across brain regions of interest [67], and the current understanding of the details of readout mechanisms remains limited. Future progress in research on readout mechanisms will be key to evaluate the different assumptions about readout in a more principled manner.

#### 3.3.2 Incorporate measurement-related considerations

When brain recordings are known to be nonlinear transformations of underlying neural activity (e.g., fMRI, in which BOLD responses are related to neural responses via the hemodynamic response function, or HRF [68], knowledge about the nonlinear relationship between the neural responses and the measurements can (and often should) be explicitly incorporated into the mapping. Failing to do so might privilege feature sets that incorporate properties of the measurement over feature sets that more accurately reflect the neural representations encoded in a brain region but happen to be nonlinearly related to the measured signal.

The issue of nonlinear properties of the measurement is prominent in fMRI analyses. The traditional approach to fMRI data fitting is a linearized encoding model: the predictor variable is convolved with the HRF, and the resulting predictor is then linearly fitted to the data. However, these models typically assume a fixed HRF across voxels [68] and/or conditions [69] which is not biologically plausible [70, 71]. More flexible finite impulse response models (FIR; [72, 73]) require fitting a large number of parameters and are computationally brittle. Thus, instead of sticking with linearized approaches, some researchers are suggesting to model the HRF shape explicitly within a family of nonlinear functions motivated by physiological data (for instance, [74, 75, 76]). By using a constrained space of nonlinear mappings (rather than an unconstrained space of linear mappings, as in FIR), one can estimate the veridical shape of the HRFs using a relatively small number of parameters. In principle, the nonlinear modeling approach can be applied to both encoding and decoding fMRI studies, although in practice the encoding direction is more feasible.

M/EEG analyses face similar issues. A common approach (for both encoding and decoding) is to fit a linear mapping between the predictors and the M/EEG measurements or their derivative (e.g., power in a particular frequency band). However, the predictor features taken from the best linear model might inadvertently incorporate the nonlinear mapping between the feature and the captured response. Further, for M/EEG, the recorded signal is a combination of both inhibitory and excitatory signals; thus, treating it as a straightforward linear combination is not always possible [77]. Thus, linear mapping models often overlook the complexities of neuroimaging signals, sacrificing biological plausibility as a result.

To summarize **Section 3**, different research goals place different constraints on the mapping model. A particular goal might require using a generic linear mapping model, adding additional restrictions to that model, using a particular class of nonlinear models, or imposing no *a priori* restrictions on the mapping space.

## 4 PRACTICAL CONSIDERATIONS

The criteria outlined above are primarily based on theoretical considerations: which mapping model has the properties that allow us to achieve a particular goal? However, another important consideration is practical feasibility: do we have enough data to accurately estimate the mapping? Will the noise in our data lead certain mapping models to fail?

Determining how much data is required for fitting a particular mapping model has critical implications for experimental design (the number of trials/data points per participant, the number of repetitions per stimulus, etc.). In general, the fewer constraints are placed on the mapping model, the more data will be needed to converge on a good mapping. This relationship can be estimated empirically using standard validation methods by, for instance, taking a large dataset and evaluating the mapping model’s predictive accuracy on left-out test data while gradually increasing the size of the training dataset. However, few studies report such analyses (and in some cases, large-enough datasets may still be lacking). One exception is a line of fMRI studies that aim to determine the best mapping model for linking interregional functional correlations and behavioral/demographic traits. The results of these studies are mixed: some report a marked advantage of nonlinear mapping models over linear ones [78] whereas others report that linear mapping models perform equally well even when the training set includes several thousand brain images [79, 80]. Thus, the field would greatly benefit from further systematic examinations of the influence of dataset size (and other experimental design properties) on the performance of a particular mapping model type.

Even with large amounts of data, certain measurement properties might force us to use a particular mapping class. For instance, Nozari et al. [81] show that the relationship between activity in different brain regions during rest, as captured by fMRI, is best modeled with linear mappings and suggest that fMRI’s inevitable spatiotemporal signal averaging might be to blame (although see [82] for contrary evidence). In sum, even after establishing theoretical desiderata for the mapping model, we need to conduct rigorous empirical tests to determine which mapping model class will achieve good predictive accuracy on test data given the measurement technique, the amount and quality of available data, and other practical considerations.

## 5 GOING FORWARD: EVALUATING MAPPING MODEL COMPLEXITY

Instead of focusing exclusively on the linear/nonlinear dichotomy, we propose to reframe the choice of mapping model in the context of a broader notion of model complexity. Complexity lies at the heart of most desiderata discussed above. Arbitrarily complex models make predictive accuracy comparisons across feature sets more difficult; they can be harder to interpret; and they are perhaps less likely to match computations in biological circuits. Thus, we suggest replacing the linear/nonlinear dichotomy with a framework that takes into account the complexity of the mapping model.

### 5.1 The role of complexity in selecting a mapping model

As we saw in **Section 3**, specific research goals impose different constraints on the mapping model. Further, these constraints are often more graded than the linear/nonlinear distinction and can instead be seen as restrictions on model complexity:

- Interpreting individual features is easier when the mapping is not only linear, but also sparse, so that each neuron can be described with only a few features. Reframing the mapping model choice in terms of complexity allows us to pick out simple mappings within the class of linear mapping models, thus facilitating interpretation.
- Satisfying biological constraints, such as accounting for physiological properties of the measurement or simulating neural readout, may require a certain degree of nonlinearity but these nonlinearities are often well-defined and can keep overall model complexity relatively low.
- Testing whether a feature set accurately captures the representational space of neural responses may require the mapping to preserve certain properties of that space. Here, the complexity of the mapping model depends primarily on the hypothesis being tested.
- Comparing and/or interpreting feature sets is possible even when the mapping is nonlinear, as long as we can compare the mappings using a metric that incorporates both predictive accuracy and model complexity.
- Decoding features from neural data and building accurate encoding models of the brain does not require placing any theory-based restrictions on the mapping model (although such restrictions might improve performance in practice).

**Figure 3** shows the research goals discussed above together with the mapping model types that are traditionally used to achieve these goals, as well as our proposal to shift from the linear/nonlinear dichotomy to explicit estimates of model complexity. Note that this diagram depicts theoretical, *a priori* criteria for restricting mapping model complexity; practical considerations might impose additional constraints to achieve better predictivity (see **Section 4)**.

**FIGURE 3.**
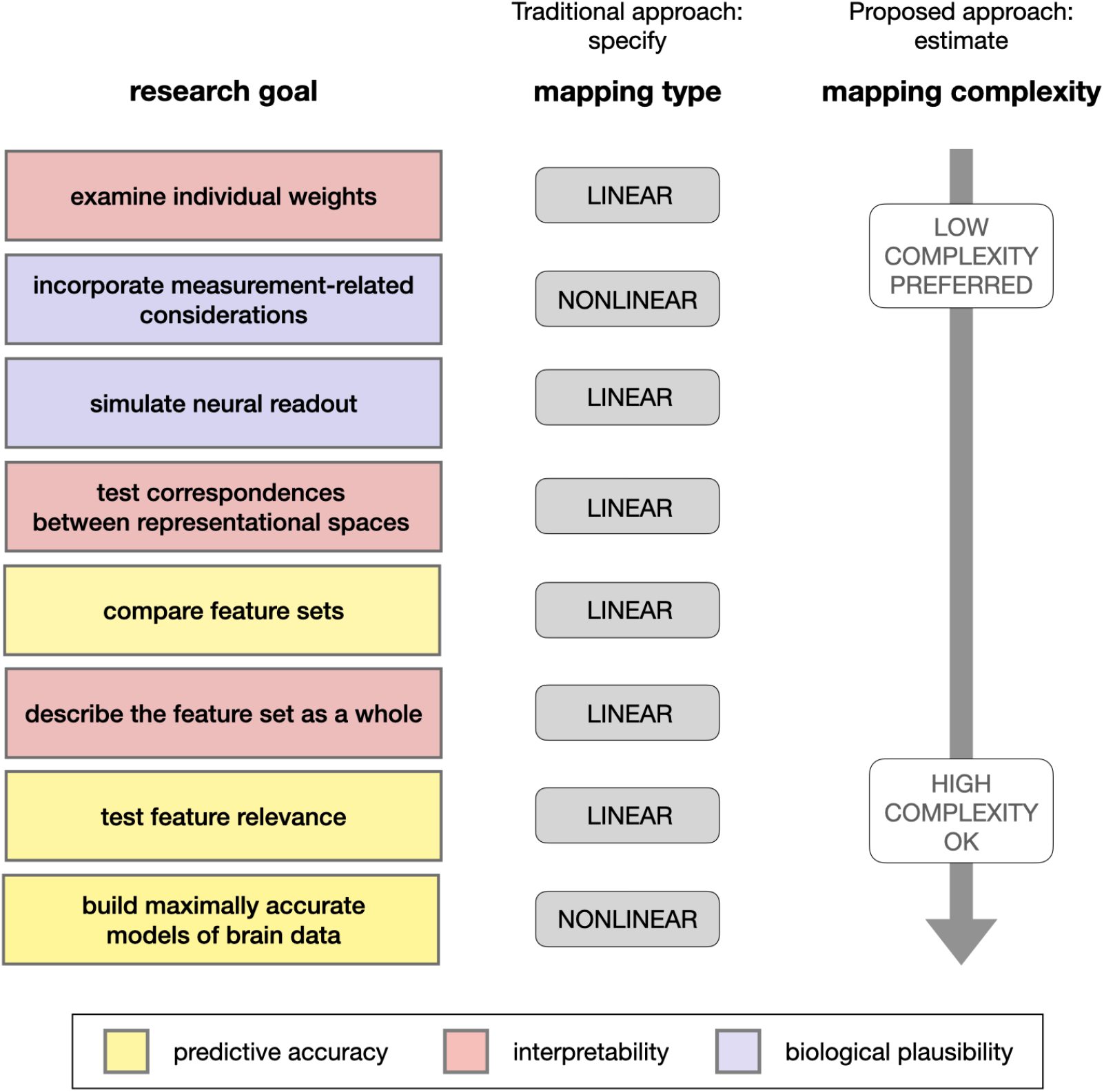
Different research goals are currently being collapsed into the “linear/nonlinear” dichotomy but in fact correspond to different degrees of mapping model complexity. Note that the exact ordering of research goals along the complexity continuum is approximate and shown primarily for illustration purposes.

### 5.2 Complexity measures

How can we estimate the complexity of mapping models? To date, many studies have focused primarily on a binary distinction in which linear models are “simple” and nonlinear models are “complex”. However, as discussed above, this distinction is overly simplistic. Here, we review several measures of mapping model complexity that are commonly used in the ML literature and may serve as an alternative to the linear/nonlinear dichotomy (see **Table 1** for a side-by-side comparison).

**TABLE 1.**
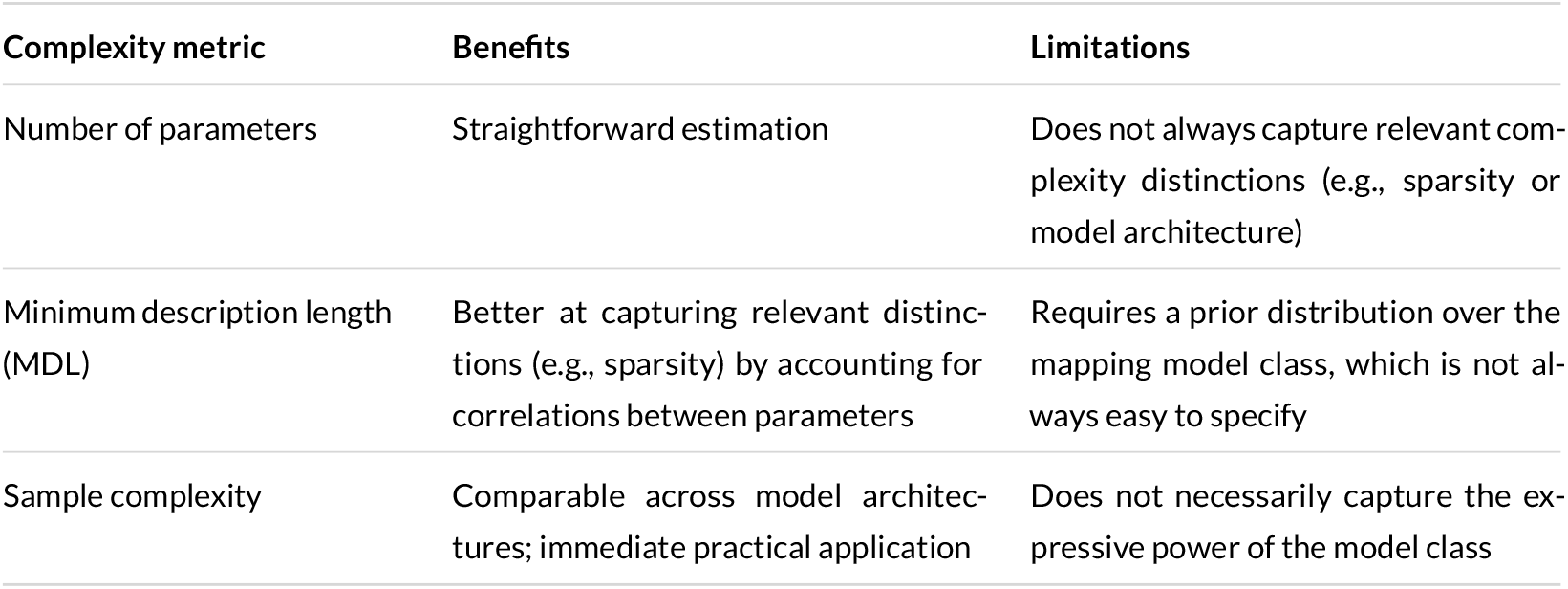
Benefits and limitations of several mapping model complexity measures.

| Complexity metric | Benefits | Limitations |
| --- | --- | --- |
| Number of parameters | Straightforward estimation | Does not always capture relevant complexity distinctions (e.g., sparsity or model architecture) |
| Minimum description length (MDL) | Better at capturing relevant distinctions (e.g., sparsity) by accounting for correlations between parameters | Requires a prior distribution over the mapping model class, which is not always easy to specify |
| Sample complexity | Comparable across model architectures; immediate practical application | Does not necessarily capture the expressive power of the model class |

#### 5.2.1 Number of free parameters

A common approach to measuring model complexity is by considering the number of free parameters in the model. In this approach, each model class encounters a penalty that corresponds to its number of parameters, such that classes with more parameters have a larger penalty. In order to justify the use of additional parameters, a model needs to achieve a substantial performance improvement compared to models with fewer parameters. This tradeoff is often implemented using Akaike’s Information Criterion (AIC) or the Bayesian Information Criterion (BIC), which reward models for good predictive performance but penalize them for the number of parameters. Although simple to estimate, this complexity measure often fails to capture distinctions that seem intuitively important. For instance, a linear and a nonlinear model with the same number of parameters would have equal complexity in this view, even though the latter often has a greater expressive power. Another example is a sparse mapping model that allows non-zero weights only for a few features vs. a dense model that places non-zero weights on, say, 500 features: if the initial feature vector size is the same, then these models will have the same number of parameters and therefore equal complexity under this measure.

#### 5.2.2 Minimum description length

Another common approach to measuring model complexity is based on minimum description length (MDL; [83]). This approach typically assumes an encoding function, i.e., a formal description language, over a class of models, and the complexity of each model within the class is determined by the length of the model’s encoding. The encoding function essentially serves as a prior over the model class: more probable mapping models would be assigned shorter descriptions (see [84, 8] for a discussion of the relationship between priors and regularization constraints). The MDL approach can overcome some of the limitations of complexity measures based solely on the number of free parameters by exploiting correlations between parameters to achieve a shorter description length. For instance, under this scheme, sparse models can have a shorter description length and would therefore be considered less complex. However, the main limitation of an MDL-based metric is that it requires specifying a mapping model class, as well as an encoding scheme for mapping models within that class. Thus, if there is no natural prior over the set of mappings we wish to compare, an architecture-free complexity measure may be preferred^4^.

#### 5.2.3 Sample complexity

Finally, a more practice-oriented metric is sample complexity [87]. Loosely speaking, the sample complexity of a model class is a function that determines the minimal number of training samples needed to achieve desired model performance. It is not always straightforward to compute this function *a priori*; however, it can be assessed empirically by computing learning curves, i.e., the achieved level of predictive accuracy on a test set as a function of the number of training samples. Estimates of sample complexity are vital for understanding whether a given model failed because the underlying hypothesis was wrong or because the dataset was too small to achieve a proper fitting.

In summary, instead of defaulting to linear models, we propose to incorporate a measure of model complexity into the general evaluation framework of encoding/decoding models. This measure can be used in different ways depending on the research goal. For instance, for feature comparison, if two feature sets produce equally accurate mapping models, the feature set corresponding to a simpler mapping model (as measured with minimum description length) may represent a better fit to neural data. For estimates of potential downstream readout, instead of limiting ourselves to linear functions, we can consider a range of possible mappings, where simpler mappings reflect a higher probability that these features are used downstream. Thus, measuring model complexity can serve as a powerful tool for mapping model evaluation and selection.

## 6 CONCLUSION

The encoding/decoding framework in contemporary cognitive neuroscience has provided many valuable insights. However, in some cases, the field has been held back by its excessive reliance on linear mappings between features and brain activity. Here, we have described various research goals that are typically considered when specifying a mapping model. Contrary to popular belief, few of these goals require the use of linear mapping models. Instead, some do not require placing any constraints on the mapping model, some require placing specific nonlinear constraints, and some use linearity simply as a proxy for reducing model complexity. We therefore suggest to explicitly include measures of model complexity when selecting and evaluating mapping models. An increased focus on mapping model complexity could help the field discover a richer space of accurate, simple, biologically plausible predictors of neural activity, thus advancing our overall understanding of brain function.

## 7 ACKNOWLEDGEMENTS

This paper is part of the Generative Adversarial Collaboration (GAC) initiative organized by the Computational Cognitive Conference board. We thank the GAC organizers, especially Megan Peters and Gunnar Blohm, for their invaluable help with this initiative. Many of the ideas discussed in this work arose during the GAC workshop in October 2020 (https://www.youtube.com/watch?v=UI5KclR71IE&list=PLNWftEg2R4s5iObSUvPXhnDJvyNbs4PnM&index=2). We thank the invited workshop speakers — Kohitij Kar, Mariya Toneva, Laura Gwilliams, Jean-Rémi King, Martin Hebart, and Anna Schapiro—as well as workshop participants for their ideas, comments, questions, and suggestions. We also thank the reviewers who provided comments on our GAC proposal (the reviews are available at https://openreview.net/forum?id=-o0dOwashib) and on earlier versions of this paper.

## Footnotes

1 Note that the neural data being fitted is not necessarily the neural recording itself: researchers may choose to predict the average firing rate, power in a particular frequency band, or beta coefficients from the general linear model (GLM) of fMRI responses [1].

2 In this paper, we discuss both encoding mapping models, i.e. models that map from the features of interest to the neural variable, and decoding mapping models, i.e. models that map from the neural variable to the features of interest (**Figure 1A**). Others have discussed the relative merits of the two approaches [2, 1, 3, 4]; our arguments in this paper apply to both mapping directions, unless specified otherwise.

3 These desiderata are commonly used to argue in favor of or against certain mapping model choices. See, e.g., the online discussion of the proposal that served as a precursor to this paper: https://openreview.net/forum?id=-o0dOwashib

4 For example, certain informational measures [e.g., 85, 86] can be used to measure the complexity of the statistical relationship between the inputs and outputs of the mapping model (e.g., features and predicted neural data) regardless of a particular architecture or model class and, in some cases, may also capture the complexity of non-parametric generative models.

## References

[1] King JR, Gwilliams L, Holdgraf C, Sassenhagen J, Barachant A, Engemann D, et al. Encoding and Decoding Framework to Uncover the Algorithms of Cognition. In: The Cognitive Neurosciences, 6th edition, vol. 6 MIT Press; 2020.p. 691–702. https://hal.archives-ouvertes.fr/hal-01848442.

[2] Holdgraf CR, Rieger JW, Micheli C, Martin S, Knight RT, Theunissen FE. Encoding and Decoding Models in Cognitive Electrophysiology. Frontiers in Systems Neuroscience 2017;11. https://www.frontiersin.org/articles/10.3389/fnsys.2017.00061/full#B83.

[3] Kriegeskorte N, Douglas P. Interpreting encoding and decoding models. Current Opinion in Neurobiology 2019;55:167–179. https://www.sciencedirect.com/science/article/pii/S0959438818301004.

[4] Naselaris T, Kay KN, Nishimoto S, Gallant JL. Encoding and decoding in fMRI. NeuroImage 2011 May;56(2):400–410.

[5] Toneva M, Wehbe L. Interpreting and improving natural-language processing (in machines) with natural language-processing (in the brain). arXiv:190511833 [cs, q-bio] 2019 Nov;http://arxiv.org/abs/1905.11833, arXiv: 1905.11833.

[6] Aertsen AMHJ, Johannesma PIM. The Spectro-Temporal Receptive Field. Biological Cybernetics 1981 Nov;42(2):133–143. https://doi.org/10.1007/BF00336731.

[7] van Gerven MAJ. A primer on encoding models in sensory neuroscience. Journal of Mathematical Psychology 2017 Feb;76:172–183. https://www.sciencedirect.com/science/article/pii/S0022249616300487.

[8] Wu MCK, David SV, Gallant JL. Complete functional characterization of sensory neurons by system identification. Annual Review of Neuroscience 2006 Jun;29(1):477–505. https://www.annualreviews.org/doi/10.1146/annurev.neuro.29.051605.113024.

[9] Bzdok D, Varoquaux G, Thirion B. Neuroimaging Research: From Null-Hypothesis Falsification to Out-of-Sample Generalization. Educational and Psychological Measurement 2017 Oct;77(5):868–880. https://doi.org/10.1177/0013164416667982.

[10] Chang N, Pyles JA, Marcus A, Gupta A, Tarr MJ, Aminoff EM. BOLD5000, a public fMRI dataset while viewing 5000 visual images. Scientific Data 2019 May;6(1):49. https://www.nature.com/articles/s41597-019-0052-3.

[11] Majaj NJ, Hong H, Solomon EA, DiCarlo JJ. Simple Learned Weighted Sums of Inferior Temporal Neuronal Firing Rates Accurately Predict Human Core Object Recognition Performance. Journal of Neuroscience 2015 Sep;35(39):13402–13418. https://www.jneurosci.org/content/35/39/13402.

[12] Schoffelen JM, Oostenveld R, Lam NHL, Uddén J, Hultén A, Hagoort P. A 204-subject multimodal neuroimaging dataset to study language processing. Scientific Data 2019 Apr;6(1):17. https://www.nature.com/articles/s41597-019-0020-y.

[13] Hasanzadeh F, Mohebbi M, Rostami R. Prediction of rTMS treatment response in major depressive disorder using machine learning techniques and nonlinear features of EEG signal. Journal of Affective Disorders 2019 Sep;256:132–142. http://www.sciencedirect.com/science/article/pii/S0165032719305543.

[14] Kazemi Y, Houghten S. A deep learning pipeline to classify different stages of Alzheimer’s disease from fMRI data. In: 2018 IEEE Conference on Computational Intelligence in Bioinformatics and Computational Biology (CIBCB); 2018. p. 1–8.

[15] Kim J, Calhoun VD, Shim E, Lee JH. Deep neural network with weight sparsity control and pre-training extracts hierarchical features and enhances classification performance: Evidence from whole-brain resting-state functional connectivity patterns of schizophrenia. NeuroImage 2016 Jan;124:127–146. http://www.sciencedirect.com/science/article/pii/S1053811915003985.

[16] Leming M, Górriz JM, Suckling J. Ensemble Deep Learning on Large, Mixed-Site fMRI Datasets in Autism and Other Tasks. International Journal of Neural Systems 2020 Jul;30(07):2050012. http://arxiv.org/abs/2002.07874, arXiv: 2002.07874.

[17] Kumar S, Yoo K, Rosenberg MD, Scheinost D, Constable RT, Zhang S, et al. An information network flow approach for measuring functional connectivity and predicting behavior. Brain and Behavior 2019;9(8):e01346. https://onlinelibrary.wiley.com/doi/abs/10.1002/brb3.1346, _eprint: https://onlinelibrary.wiley.com/doi/pdf/10.1002/brb3.1346.

[18] Morioka H, Calhoun V, Hyvärinen A. Nonlinear ICA of fMRI reveals primitive temporal structures linked to rest, task, and behavioral traits. NeuroImage 2020 Sep;218:116989. http://www.sciencedirect.com/science/article/pii/S1053811920304754.

[19] Xiao L, Stephen JM, Wilson TW, Calhoun VD, Wang YP. Alternating Diffusion Map Based Fusion of Multimodal Brain Connectivity Networks for IQ Prediction. IEEE Transactions on Biomedical Engineering 2019 Aug;66(8):2140–2151. Conference Name: IEEE Transactions on Biomedical Engineering.

[20] Caucheteux C, King JR. Language processing in brains and deep neural networks: computational convergence and its limits. bioRxiv 2020 Jul;p. 2020.07.03.186288. https://www.biorxiv.org/content/10.1101/2020.07.03.186288v1.

[21] Jain S, Vo VA, Mahto S, LeBel A, Turek JS, Huth AG. Interpretable multi-timescale models for predicting fMRI responses to continuous natural speech. bioRxiv 2020 Oct;p. 2020.10.02.324392. https://www.biorxiv.org/content/10.1101/2020.10.02.324392v1.

[22] Schrimpf M, Kubilius J, Hong H, Majaj NJ, Rajalingham R, Issa EB, et al. Brain-Score: Which Artificial Neural Network for Object Recognition is most Brain-Like? bioRxiv 2018 Sep;p. 407007. https://www.biorxiv.org/content/10.1101/407007v1.

[23] Yamins DLK, Hong H, Cadieu CF, Solomon EA, Seibert D, DiCarlo JJ. Performance-optimized hierarchical models predict neural responses in higher visual cortex. Proceedings of the National Academy of Sciences of the United States of America 2014 Jun;111(23):8619–8624.

[24] Anderson AJ, Binder JR, Fernandino L, Humphries CJ, Conant LL, Aguilar M, et al. Predicting Neural Activity Patterns Associated with Sentences Using a Neurobiologically Motivated Model of Semantic Representation. Cerebral Cortex 2017 Sep;27(9):4379–4395. http://academic.oup.com/cercor/article/27/9/4379/3056416.

[25] Lee Masson H, Isik L. Functional selectivity for naturalistic social interaction perception in the human superior temporal sulcus. bioRxiv 2021 Mar;p. 2021.03.26.437258. https://www.biorxiv.org/content/10.1101/2021.03.26.437258v1.

[26] Sudre G, Pomerleau D, Palatucci M, Wehbe L, Fyshe A, Salmelin R, et al. Tracking neural coding of perceptual and semantic features of concrete nouns. NeuroImage 2012 Aug;62(1):451–463. https://www.sciencedirect.com/science/article/pii/S1053811912004442.

[27] Haufe S, Meinecke F, Görgen K, Dähne S, Haynes JD, Blankertz B, et al. On the interpretation of weight vectors of linear models in multivariate neuroimaging. NeuroImage 2014 Feb;87:96–110.

[28] Kamitani Y, Tong F. Decoding the visual and subjective contents of the human brain. Nature neuroscience 2005 May;8(5):679–685. https://www.ncbi.nlm.nih.gov/pmc/articles/PMC1808230/.

[29] Kriegeskorte N. Pattern-information analysis: From stimulus decoding to computational-model testing. NeuroImage 2011 May;56(2):411–421. http://www.sciencedirect.com/science/article/pii/S1053811911000978.

[30] Kay KN. Principles for models of neural information processing. NeuroImage 2018 Oct;180(Pt A):101–109.

[31] Kording KP, Blohm G, Schrater P, Kay K. Appreciating the variety of goals in computational neuroscience. Neurons, Behavior, Data analysis, and Theory 2020 Feb;3(6). http://arxiv.org/abs/2002.03211, arXiv: 2002.03211.

[32] Kanwisher N, McDermott J, Chun MM. The Fusiform Face Area: A Module in Human Extrastriate Cortex Specialized for Face Perception. Journal of Neuroscience 1997 Jun;17(11):4302–4311. https://www.jneurosci.org/content/17/11/4302.

[33] Cadieu CF, Hong H, Yamins DLK, Pinto N, Ardila D, Solomon EA, et al. Deep Neural Networks Rival the Representation of Primate IT Cortex for Core Visual Object Recognition. PLOS Computational Biology 2014 Dec;10(12):e1003963. https://journals.plos.org/ploscompbiol/article?id=10.1371/journal.pcbi.1003963.

[34] Dapello J, Marques T, Schrimpf M, Geiger F, Cox D, DiCarlo JJ. Simulating a Primary Visual Cortex at the Front of CNNs Improves Robustness to Image Perturbations. In: Larochelle H, Ranzato M, Hadsell R, Balcan MF, Lin H, editors. Advances in Neural Information Processing Systems, vol. 33 Curran Associates, Inc.; 2020. p. 13073–13087. https://proceedings.neurips.cc/paper/2020/file/98b17f068d5d9b7668e19fb8ae470841-Paper.pdf.

[35] David SV, Gallant JL. Predicting neuronal responses during natural vision. Network (Bristol, England) 2005 Sep;16(2-3):239–260.

[36] Geiger F, Schrimpf M, Marques T, DiCarlo JJ. Wiring Up Vision: Minimizing Supervised Synaptic Updates Needed to Produce a Primate Ventral Stream. bioRxiv; 2020.

[37] Hsu A, Borst A, Theunissen FE. Quantifying variability in neural responses and its application for the validation of model predictions. Network (Bristol, England) 2004 May;15(2):91–109.

[38] Lage-Castellanos A, Valente G, Formisano E, Martino FD. Methods for computing the maximum performance of computational models of fMRI responses. PLOS Computational Biology 2019 Mar;15(3):e1006397. https://journals.plos.org/ploscompbiol/article?id=10.1371/journal.pcbi.1006397.

[39] Schoppe O, Harper NS, Willmore BDB, King AJ, Schnupp JWH. Measuring the Performance of Neural Models. Frontiers in Computational Neuroscience 2016;10. https://www.frontiersin.org/article/10.3389/fncom.2016.00010.

[40] Schrimpf M, Blank I, Tuckute G, Kauf C, Hosseini EA, Kanwisher N, et al. Artificial Neural Networks Accurately Predict Language Processing in the Brain. bioRxiv 2020 Jun;p. 2020.06.26.174482. https://www.biorxiv.org/content/10.1101/2020.06.26.174482v1.

[41] Talebi V, Baker CL. Natural versus Synthetic Stimuli for Estimating Receptive Field Models: A Comparison of Predictive Robustness. Journal of Neuroscience 2012 Feb;32(5):1560–1576. https://www.jneurosci.org/content/32/5/1560.

[42] Naselaris T, Olman CA, Stansbury DE, Ugurbil K, Gallant JL. A voxel-wise encoding model for early visual areas decodes mental images of remembered scenes. NeuroImage 2015 Jan;105:215–228.

[43] Schrimpf M, Kubilius J, Lee MJ, Murty NAR, Ajemian R, DiCarlo JJ. Integrative Benchmarking to Advance Neurally Mechanistic Models of Human Intelligence. Neuron 2020 Nov;108(3):413–423. https://www.cell.com/neuron/abstract/S0896-6273(20)30605-X.

[44] Khosla M, Wehbe L. High-level visual areas act like domain-general filters with strong selectivity and functional specialization. bioRxiv 2022;.

[45] Ratan Murty NA, Bashivan P, Abate A, DiCarlo JJ, Kanwisher N. Computational models of category-selective brain regions enable high-throughput tests of selectivity. Nature Communications 2021 Sep;12(1):5540. https://www.nature.com/articles/s41467-021-25409-6, bandiera_abtest: a Cc_license_type: cc_by Cg_type: Nature Research Journals Number: 1 Primary_atype: Research.

[46] Desimone R, Albright TD, Gross CG, Bruce C. Stimulus-selective properties of inferior temporal neurons in the macaque. Journal of Neuroscience 1984 Aug;4(8):2051–2062. https://www.jneurosci.org/content/4/8/2051.

[47] Buzsáki G. The brain from inside out. Oxford University Press; 2019.

[48] Ponce CR, Xiao W, Schade PF, Hartmann TS, Kreiman G, Livingstone MS. Evolving Images for Visual Neurons Using a Deep Generative Network Reveals Coding Principles and Neuronal Preferences. Cell 2019 May;177(4):999–1009.e10. https://www.sciencedirect.com/science/article/pii/S0092867419303915.

[49] Kay KN, Naselaris T, Prenger RJ, Gallant JL. Identifying natural images from human brain activity. Nature 2008 Mar;452(7185):352–355. https://www.ncbi.nlm.nih.gov/pmc/articles/PMC3556484/.

[50] Jain S, Huth AG. Incorporating Context into Language Encoding Models for fMRI. bioRxiv 2018 Nov;p. 327601. https://www.biorxiv.org/content/10.1101/327601v2.

[51] Richards BA, Lillicrap TP, Beaudoin P, Bengio Y, Bogacz R, Christensen A, et al. A deep learning framework for neuroscience. Nature Neuroscience 2019 Nov;22(11):1761–1770. https://www.nature.com/articles/s41593-019-0520-2, number: 11.

[52] Chaudhuri R, Gerçek B, Pandey B, Peyrache A, Fiete I. The intrinsic attractor manifold and population dynamics of a canonical cognitive circuit across waking and sleep. Nature Neuroscience 2019 Sep;22(9):1512–1520. https://www.nature.com/articles/s41593-019-0460-x, bandiera_abtest: a Cg_type: Nature Research Journals Number: 9 Primary_atype: Research.

[53] Gallego JA, Perich MG, Naufel SN, Ethier C, Solla SA, Miller LE. Cortical population activity within a preserved neural manifold underlies multiple motor behaviors. Nature Communications 2018 Oct;9(1):4233. https://www.nature.com/articles/s41467-018-06560-z, bandiera_abtest: a Cc_license_type: cc_by Cg_type: Nature Research Journals Number: 1 Primary_atype: Research.

[54] Jazayeri M, Ostojic S. Interpreting neural computations by examining intrinsic and embedding dimensionality of neural activity. arXiv:210704084 [q-bio] 2021 Aug; http://arxiv.org/abs/2107.04084, arXiv: 2107.04084.

[55] Barak O, Rigotti M, Fusi S. The Sparseness of Mixed Selectivity Neurons Controls the Generalization–Discrimination Trade-Off. Journal of Neuroscience 2013 Feb;33(9):3844–3856. https://www.jneurosci.org/content/33/9/3844.

[56] Barlow H. Trigger features, adaptation and economy of impulses. In: Information Processing in the Nervous System Springer; 1969.p. 209–230.

[57] Olshausen BA, Field DJ. Sparse coding of sensory inputs. Current Opinion in Neurobiology 2004 Aug;14(4):481–487. https://www.sciencedirect.com/science/article/pii/S0959438804001035.

[58] Vinje WE, Gallant JL. Sparse Coding and Decorrelation in Primary Visual Cortex During Natural Vision. Science 2000 Feb;287(5456):1273–1276. https://science.sciencemag.org/content/287/5456/1273.

[59] Beniaguev D, Segev I, London M. Single cortical neurons as deep artificial neural networks. Neuron 2021 Sep;109(17):2727–2739.e3. https://www.cell.com/neuron/abstract/S0896-6273(21)00501-8.

[60] Ghazanfar AA, Nicolelis MA. Nonlinear processing of tactile information in the thalamocortical loop. Journal of Neurophysiology 1997 Jul;78(1):506–510.

[61] Gidon A, Zolnik TA, Fidzinski P, Bolduan F, Papoutsi A, Poirazi P, et al. Dendritic action potentials and computation in human layer 2/3 cortical neurons. Science 2020 Jan;367(6473):83–87. https://science.sciencemag.org/content/367/6473/83.

[62] Jones IS, Kording KP. Might a Single Neuron Solve Interesting Machine Learning Problems Through Successive Computations on Its Dendritic Tree? Neural Computation 2021 May;33(6):1554–1571. https://doi.org/10.1162/neco_a_01390.

[63] Shamir M, Sompolinsky H. Nonlinear Population Codes. Neural Computation 2004 Jun;16(6):1105–1136. https://doi.org/10.1162/089976604773717559.

[64] Ritchie JB, Kaplan DM, Klein C. Decoding the Brain: Neural Representation and the Limits of Multivariate Pattern Analysis in Cognitive Neuroscience. The British Journal for the Philosophy of Science 2019 Jun;70(2):581–607. https://doi.org/10.1093/bjps/axx023.

[65] Kukačka J, Golkov V, Cremers D. Regularization for Deep Learning: A Taxonomy. arXiv:171010686 [cs, stat] 2017 Oct; http://arxiv.org/abs/1710.10686, arXiv: 1710.10686.

[66] Yoshida T, Ohki K. Natural images are reliably represented by sparse and variable populations of neurons in visual cortex. Nature Communications 2020 Feb;11(1):872. https://www.nature.com/articles/s41467-020-14645-x.

[67] Anzellotti S, Coutanche MN. Beyond Functional Connectivity: Investigating Networks of Multivariate Representations. Trends in Cognitive Sciences 2018 Mar;22(3):258–269. https://www.sciencedirect.com/science/article/pii/S1364661317302620.

[68] Friston KJ, Mechelli A, Turner R, Price CJ. Nonlinear responses in fMRI: the Balloon model, Volterra kernels, and other hemodynamics. NeuroImage 2000 Oct;12(4):466–477.

[69] Pedregosa F, Eickenberg M, Ciuciu P, Thirion B, Gramfort A. Data-driven HRF estimation for encoding and decoding models. NeuroImage 2015 Jan;104:209–220. https://www.sciencedirect.com/science/article/pii/S1053811914008027.

[70] Ekstrom AD. Regional variation in neurovascular coupling and why we still lack a Rosetta Stone. Philosophical Transactions of the Royal Society B: Biological Sciences 2021 Jan;376(1815):20190634. https://royalsocietypublishing.org/doi/abs/10.1098/rstb.2019.0634.

[71] Handwerker DA, Ollinger JM, D’Esposito M. Variation of BOLD hemodynamic responses across subjects and brain regions and their effects on statistical analyses. NeuroImage 2004 Apr;21(4):1639–1651.

[72] Dale AM. Optimal experimental design for event-related fMRI. Human Brain Mapping 1999;8(2-3):109–114. https://onlinelibrary.wiley.com/doi/abs/10.1002/%28SICI%291097-0193%281999%298%3A2/3%3C109%3A%3AAID-HBM7%3E3.0.CO%3B2-W, _eprint: https://onlinelibrary.wiley.com/doi/pdf/10.1002/%28SICI%291097-0193%281999%298%3A2/3%3C109%3A%3AAID-HBM7%3E3.0.CO%3B2-W.

[73] Glover GH. Deconvolution of Impulse Response in Event-Related BOLD fMRI1. NeuroImage 1999 Apr;9(4):416–429. https://www.sciencedirect.com/science/article/pii/S1053811998904190.

[74] Lindquist MA, Wager TD. Validity and power in hemodynamic response modeling: a comparison study and a new approach. Human Brain Mapping 2007 Aug;28(8):764–784.

[75] Shain C, Blank IA, van Schijndel M, Schuler W, Fedorenko E. fMRI reveals language-specific predictive coding during naturalistic sentence comprehension. Neuropsychologia 2020;138:107307.

[76] Shain C. CDRNN: Discovering Complex Dynamics in Human Language Processing. In: Proceedings of the 59th Annual Meeting of the Association for Computational Linguistics and the 11th International Joint Conference on Natural Language Processing (Volume 1: Long Papers) Online: Association for Computational Linguistics; 2021. p. 3718–3734. https://aclanthology.org/2021.acl-long.288.

[77] Hansen PC, Kringelbach ML, Salmelin R, editors. MEG: An introduction to methods. MEG: An introduction to methods, New York, NY, US: Oxford University Press; 2010. Pages: xii, 436.

[78] Bertolero MA, Bassett DS. Deep Neural Networks Carve the Brain at its Joints. arXiv:200208891 [physics, q-bio] 2020 Feb; http://arxiv.org/abs/2002.08891, arXiv: 2002.08891.

[79] He T, Kong R, Holmes AJ, Nguyen M, Sabuncu MR, Eickhoff SB, et al. Deep neural networks and kernel regression achieve comparable accuracies for functional connectivity prediction of behavior and demographics. NeuroImage 2020 Feb;206:116276. https://www.sciencedirect.com/science/article/pii/S1053811919308675.

[80] Schulz MA, Yeo BTT, Vogelstein JT, Mourao-Miranada J, Kather JN, Kording K, et al. Different scaling of linear models and deep learning in UKBiobank brain images versus machine-learning datasets. Nature Communications 2020 Aug;11(1):4238. https://www.nature.com/articles/s41467-020-18037-z.

[81] Nozari E, Stiso J, Caciagli L, Cornblath EJ, He X, Bertolero MA, et al. Is the brain macroscopically linear? A system identification of resting state dynamics. arXiv:201212351 [cs, eess, math, q-bio] 2020 Dec;http://arxiv.org/abs/2012.12351, arXiv: 2012.12351.

[82] Anzellotti S, Fedorenko E, Kell AJE, Caramazza A, Saxe R. Measuring and Modeling Nonlinear Interactions Between Brain Regions with fMRI. bioRxiv 2017 Sep;p. 074856. https://www.biorxiv.org/content/10.1101/074856v2.

[83] Rissanen J. Modeling by shortest data description. Automatica 1978 Sep;14(5):465–471. https://www.sciencedirect.com/science/article/pii/0005109878900055.

[84] Diedrichsen J, Kriegeskorte N. Representational models: A common framework for understanding encoding, pattern-component, and representational-similarity analysis. PLOS Computational Biology 2017 Apr;13(4):e1005508. https://journals.plos.org/ploscompbiol/article?id=10.1371/journal.pcbi.1005508.

[85] Bialek W, Nemenman I, Tishby N. Predictability, complexity, and learning. Neural Computation 2001 Nov;13(11):2409–2463.

[86] Gilad-Bachrach R, Navot A, Tishby N. An Information Theoretic Tradeoff between Complexity and Accuracy. In: Schölkopf B, Warmuth MK, editors. Learning Theory and Kernel Machines Lecture Notes in Computer Science, Berlin, Heidelberg: Springer; 2003. p. 595–609.

[87] Kearns MJ, Vazirani U. An Introduction to Computational Learning Theory. MIT Press; 1994. http://direct.mit.edu/books/book/2604/An-Introduction-to-Computational-Learning-Theory.

